## Supplementary File 3 for "Analysis of zebrafish periderm enhancers facilitates identification of a regulatory variant near human *KRT8/18*"

**a Summary of ClustalO alignment between zebrafish *ppl*-10 enhancers and PPL-8.3**

| **Sequence Two (pairwise), or Sequence Three (3-way)** | **Length (bp)** | **# of identical positions in alignment (*)** | **# of null characters inserted (-)** | **% identity of human 400 bp** | **% identity, normalized by insertions** | **5-mers** | **6-mers** | **7-mers** | **8-mers** | **9-mers** | **10-mers** | **11-mers** |
| --- | --- | --- | --- | --- | --- | --- | --- | --- | --- | --- | --- | --- |
| Mouse plus strand (+) | 409 | 231 | 57 | 57.8% | 50.5% | 10 | 0 | 2 | 2 | 2 | 2 | 1 |
| Mouse minus strand (-) | 409 | 159 | 117 | 39.8% | 30.8% | 2 | 2 | 1 | 0 | 1 | 0 | 0 |
| Mouse reverse sequence | 409 | 167 | 85 | 41.8% | 34.4% | 2 | 0 | 0 | 0 | 0 | 0 | 0 |
| Mouse scrambled sequence 1 | 409 | 157 | 71 | 39.3% | 33.3% | 1 | 0 | 0 | 0 | 0 | 0 | 0 |
| Mouse scrambled sequence 2 | 409 | 163 | 81 | 40.8% | 33.9% | 3 | 1 | 0 | 0 | 0 | 0 | 0 |
| Mouse scrambled sequence 3 | 409 | 160 | 59 | 40.0% | 34.9% | 1 | 1 | 0 | 0 | 0 | 0 | 0 |
| Zebrafish plus strand (+) | 467 | 181 | 73 | 45.3% | 38.3% | 5 | 0 | 0 | 0 | 0 | 0 | 0 |
| Zebrafish minus strand (-) | 467 | 158 | 107 | 39.5% | 31.2% | 2 | 0 | 0 | 0 | 0 | 0 | 0 |
| Zebrafish reverse sequence | 467 | 168 | 107 | 42.0% | 33.1% | 2 | 0 | 1 | 0 | 0 | 0 | 0 |
| Zebrafish scrambled sequence 1 | 467 | 168 | 95 | 42.0% | 33.9% | 4 | 1 | 0 | 0 | 0 | 0 | 0 |
| Zebrafish scrambled sequence 2 | 467 | 166 | 135 | 41.5% | 31.0% | 2 | 1 | 0 | 0 | 0 | 0 | 0 |
| Zebrafish scrambled sequence 3 | 467 | 174 | 73 | 43.5% | 36.8% | 1 | 2 | 1 | 0 | 0 | 0 | 0 |
| 3-way w/ zf plus strand (+) | 467 | 96 | 158 | 24.0% | 17.2% | 0 | 0 | 0 | 0 | 0 | 0 | 0 |
| 3-way w/ zf minus strand (-) | 467 | 97 | 227 | 24.3% | 15.5% | 0 | 0 | 0 | 0 | 0 | 0 | 0 |
| 3-way w/ zf reverse sequence | 467 | 99 | 248 | 24.8% | 15.3% | 0 | 0 | 0 | 0 | 0 | 0 | 0 |
| 3-way w/ zf scrambled sequence 1 | 467 | 110 | 170 | 27.5% | 19.3% | 1 | 1 | 0 | 0 | 0 | 0 | 0 |
| 3-way w/ zf scrambled sequence 2 | 467 | 86 | 216 | 21.5% | 14.0% | 0 | 0 | 0 | 0 | 0 | 0 | 0 |
| 3-way w/ zf scrambled sequence 3 | 467 | 90 | 140 | 22.5% | 16.7% | 0 | 0 | 0 | 0 | 0 | 0 | 0 |

**b Details for ClustalO alignment between zebrafish *ppl*-10 enhancers and PPL-8.3**

**Alignment #1.** Shown below is the pair-wise alignment of the core human *periplakin* (*ppl*) enhancer conserved in mammals (400 bp) aligned to the homologous 409 bp block in mouse. These sequences correspond to the plus strand (+) sequences located upstream of *ppl*, which is transcribed to the left in each genome. Alignment #1 serves as a positive control for the described enhancer homology tests.

```
                                                                      9-mer (a)
Hu_400+      TTCTGACACAACCTGCTACACAT----CTTGGATCCCACTTGTCAAGCCTGGCCCTAGGC	56
Mm_409+      CTATCCCACAGACTGGTGACCCTGGAGCTCAGACACTCCACTGGAGAGCTATCCCTGGGC	60
              * *  ****  *** *   * *    **  **  *  *     *   **  **** ***
                                            10-mer (b)    10-mer (c)
Hu_400+      CTCTCAGG------GGTGGGTACATCAAGTGAGGAAGTCACACTGTCAGGGCAGAAAGAG	110
Mm_409+      CTCTCAAATTCACCATGAGAGCTTCTCTGCAAGGAAGTCACCCCATCAGGGCAGA--GTG	118
             ******            *         *  ********** *  **********  * *
                   8-mer (d)  9-mer (e)            (f)   (g)       7-mer (h)
Hu_400+      GGGCTGGGTGGGGCCGGACAGGCTCTGC-TGGCTCAGGATCCCGGCAGGACAGGCTGCCC	169
Mm_409+      GTTCTAGGTGGGGCTAAACAGGCTCTACCAGGCACAGTATCCCTGCAGGGCAGACTGCCC	178
             *  ** ********   ********* *  *** *** ***** ***** *** ******
                                                      (i)        (j)
Hu_400+      AACCCAGCCCCAAAACTCCCATCTTCTGTCCAACGCCCAAGCCTGCTTCCTCCCTGCCCC	229
Mm_409+      AGCCTCCTGTTCACG--------------CCCATGCTTTTTCCTGCCTGCTGCCTGCTCC	224
             * **        *                ** * **     ***** * ** ***** **
                                              (k)            (l)   (m)
Hu_400+      ---------------ACCTCACCTCACACTGCCCAGACTGGTGGGGCTGCATTTTTGGGG	274
Mm_409+      TCCCCTTCTGGGCCGTGCCAGGCTCATGCAGAGCAGACCAGTACAGCCGCATTCTTGGGC	284
                              *    ****  * *  *****  **   ** ***** *****
                       7-mer (n)               8-mer(o) 11-mer (p)
Hu_400+      C-CGGTGCCAGGTGTGATGGA-----GGACAGCCCAGGTCAGGAGGTCAGGGCCCCGTTT	328
Mm_409+      AAGTGTCTCTGGTGTGAATGACCCTGGGGCTGGTCAGGTCAGAAGGTCAGGGCCTTGTTC	344
                 **  * *******  **     ** * *  ******** ***********  ***
             (q)                                                 (r)
Hu_400+      CCTCCCAAGCCAGGATGACTCAGTGATGGACAGACTAGGCCACAAGGGG-CCGAACACAA	387
Mm_409+      CCTCCAGTGCCCAAGTGATA--------GATGAGTCAGGCTACAAAGGGGCAGAACAAGA	396
             *****   ***    ***          **      **** **** *** * *****  *
               (s)
Hu_400+      AGCCAAATTAGGG	400
Mm_409+      AACCAAACCGAGT	409
             * *****    *
```

**Alignment #2.** Shown below is the pair-wise alignment of the core human *ppl* enhancer (Hu_400+) aligned to the reverse complement of the homologous 409 bp block in mouse (Mm_409-).

```
Hu_400+      ------------------TTCTGACACAACCTGCTACACATCTTGGATCCCACTTGTCA-	41
Mm_409-      ACTCGGTTTGGTTTCTTGTTCTGCCCCTTTGTAGCCTGACTCATCTATCACTTGGGCACT	60
                               ***** * *    *        ** *  *** *    *

Hu_400+      ------AGCCTGGCCCTAGGCCTCTCAGGGGTGGGTACATCAAGTGAGGAAGTCACACTG	95
Mm_409-      GGAGGGAACAAGGCCCTGACCTTCTGACCTG----------ACCAGCCCCAGGGTCATTC	110
                   * *  ******   * *** *   *          *   *    **   ** *
                                                      7-mer
Hu_400+      TCAGGGCAGAAAGAGGGGCTGGGTGGGGCCGG-----ACAGGCTCTGCTGGCTCAGGATC	150
Mm_409-      ACACCAGAGACACTTGCCCAAGAATGCGGCTGTACTGGTCTGCTCTGCATGAGCCTGGCA	170
              **    *** *   *  *  *   * * * *         *******  *  *  *

Hu_400+      CCGGCAGGACAGGCTGCC----CAACCCAGCCCCAAAACTC----------------CCA	190
Mm_409-      CGGCCCAGAAGGGGAGGAGCAGGCAGCAGGCAGGAAAAAGCATGGGCGTGAACAGGAGGC	230
             * * *  **  **  *        * *  **   ****  *
                                                            9-mer
Hu_400+      TCTTCTGTCCAACGCCC---------AAGCCTGCT----TCCTCCCTGCCCCACCTCACC	237
Mm_409-      TGGGCAGTCTGCCCTGCAGGGATACTGTGCCTGGTAGAGCCTGTTTAGCCCCACCTAGAA	290
             *   * ***   *   *           ***** *     *      *********

Hu_400+      TCACACTGCCCAGACTGGTGGGGCTGCATTTTTGGGGCCGGTGCCAGGTGTGATGGAGGA	297
Mm_409-      CCACTCTGCCCTGATGGGGTG-ACTTCCTTGCAGAGAA--------GCTCTCATGGTGAA	341
              *** ****** **  **  *  ** * **   * *          * * * **** * *

Hu_400+      CAGCCCAGGTCAGGAGGTCAGGGCCCCGTTTCCTCCCAAGCCAGGATGACTCAGTGATGG	357
Mm_409-      TTTGAGAGGCCCA-----------------------------GGGATAGCTCTCCAGTGG	372
                   *** *                                ****  ***     ***

Hu_400+      ACAGACTAGGCCACAAGGGGCCGAACACAAAGCCAAATTAGGG	400
Mm_409-      AGTGTCTGAGCTCCAGGGTCACCAGTCTGTGG-GATAG-----	409
             *  * **  **  ** **   * *       *  * *
```

**Alignment #3.** Shown below is the pair-wise alignment of the core human *ppl* enhancer (Hu_400+) aligned to the reverse sequence of the homologous 409 bp block in mouse (Mm_409R). This alignment constitutes a negative control as the reverse sequence is a non-biological sequence.

```
Hu_400+      TTCTGACACAACCTGCTACACATCTTGGATCCCACTTGTCAAGCCTGGCCCTAGGCCTCT	60
Mm_409R      ---TGAGCCAAACC----------AAAGAACAAGACGGGGAAACATCG------------	35
                ***  *** *              ** *      *  ** * * *

Hu_400+      CAGGGGTGGGTACATCAA---GTGAGGAAGTCACAC-TGTCAGGGCAGAAAGAG-GGGCT	115
Mm_409R      ---GACTGAGTAGATAGTGAACCCGTGACCTCCCTTGTTCCGGGACTGGAAGACTGGACT	92
                *  ** *** **           **  ** *   *  * ** * * ****  ** **

Hu_400+      GGGTGGGGCCGGACAGGCTCTGCTGGCTCAGGA--TCCCGGCAGGACAGGCTGCCCAACC	173
Mm_409R      GGTCGGGGTCCCAGTAAGTGTGGTCTCTGTGAACGGGTTCTTACGCCGACATGACCAGAC	152
             **  **** *  *     * ** *  **  * *         * * *    ** ***  *

Hu_400+      CAGCCCC-----AAAACTCCCA-----TCTTCTGTCCAACGCCCAAGCCTGCTTCCTCCC	223
Mm_409R      GAGACGTACTCGGACCGTGCCGGGTCTTCCCCTCCTCGTCCGTCGTCCGTCCTTTTTCGT	212
              ** *        *   * **      **  **   *  *   *   * * ***  **

Hu_400+      TGCCCCACCTCACCTCACACTGCCCAGACTGGTG---------GGGCTGCATTTTTGGG-	273
Mm_409R      ACCCGCACTTGTCCTCCGACCCGTCAGACGGGACGTCCCTATGACACGGACCATCTCGGA	272
               ** *** *  ****  **    ***** **              * *    * * **

Hu_400+      --------GCCGGTGCCAGGTGTGATGGAGGACAGCCCAGGTCAGGAGGTCAGGGCCCCG	325
Mm_409R      CAAATCGGGGTGGATCTTGGTGAGACGGGACTACCCCACTGAAGGAACGTCTCTTCGAGA	332
                     *  **  *  **** ** **       **   *   * * ***    *

Hu_400+      TTTCCTCCCAAGC---CAGGATGACTCAGTGAT-------GGACAGA--CTAGGCCACAA	373
Mm_409R      GTACCACTTAAACTCTCCGGGTCCCTATCGAGAGGTCACCTCACAGACTCGAGGTCCCAG	392
              * ** *  ** *   * ** *  **                *****  * *** * **

Hu_400+      GGGGCCGAACACAAAGCCAAATTAGGG	400
Mm_409R      TGGTCAGACACCCTAT---------C-	409
              ** * **   *  *
```

**Alignments #’s 4, 5, and 6.** Shown below are three pair-wise alignments of the core human *ppl* enhancer (Hu_400+) aligned to one of three (S1, S2, and S3) different Fisher-Yates shuffled sequences of the homologous 409 bp plus-strand block in mouse (Mm_409+S1/S2/S3). These pair-wise alignments serve as negative controls.

```
Alignment #4

Hu_400+        TTCTGACACAACCTGCTACACATCTTGGATCCCACTTGTCAAGCCTGGCCCTAGGCCTCT	60
Mm_409+S1      --------AGAGCTC------CGCTTGG---CCTGTGCTCAAATG------------AAG	31
                         * **         *****   **  *  ****

Hu_400+        CAGGGGTGGGTACATCAAGTGAGGAAGTCACAC--TGTCAGGGCAGAAAGAGGGGCTGGG	118
Mm_409+S1      CAGAGACCGGCCAAGCACAAGGGGCGTTTACAACTTGCTGCGGGAAGTAGAGCTTCTGAG	91
               *** *   **   * **   * **   * ***   **    ** *   ****   *** *

Hu_400+        TGGGGCCGGACAGG---CTCTGCTGGCTCAGGATCCCGGCAGGACAGGCTGCCCAACCCA	175
Mm_409+S1      ACCGACCGGCCGGCACACCCTACTTGGCTTGAACCACCAATTTGTGGTGCACATGATCCT	151
                  * **** * *    * ** ** *    * * * *         *    *   * **

Hu_400+        GCCCCAAAACTCCCATCTTCTGTCCAACGCCCAAGCCTGCTTCCTCCCTGCCCCACCT--	233
Mm_409+S1      TCCGGACCGCACCAACGATTTGACGGAGTCCGCCTGCAGATTGGGGCCGAGCTGTTTTGG	211
                **  *   * ** *   * ** *  *  **     * * **    **   *     *

Hu_400+        ----CACCTCACACTGCCCAGACTGG---------TGGGGCTGCATTTTTGGGGCCGGTG	280
Mm_409+S1      AACCGGAGTATAAGTCCTCTGTGTCTTCGTCTCTCAGTGGTTTCGTTATGT--CCCTCCG	269
                       *   * * * * *  *            * ** * * ** *     **   *

Hu_400+        CCAGGTGTGATGGAGGACAGCCC-------AGGTCAGGAGG---TCAGGGCCCCGTTTCC	330
Mm_409+S1      CCTCACGGGATCAATGGCACCGCATCCCCAAGCACGGGTGCGAAGACAGGCAGAGATTCT	329
               **    * ***  * * ** * *       **  * ** *        ***   * ***

Hu_400+        TCCCAAGCCAGGATGACTCAGTGATGGACAGACTAGGCCACAAGGGGCCGAACACAAAGC	390
Mm_409+S1      GGCCACGTCAATACTCCCCGTAACTCCACAGTACAGCGCTAATAGTGGGTAGTCCGCCCT	389
                 *** * **  *   * *     *  ****   **  *  *  * *   *   *

Hu_400+        CAAATTAGGG----------	400
Mm_409+S1      ATACGGAGAGCCGCCCACCC	409
                 *   ** *
```

```
Alignment #5

Hu_400+        ---TTCTGACACAACCTGCTACACATCT----------TGGATCCCACTTGTCAAGCCTG	47
Mm_409+S2      TAAACGAGGAGCTACCTCATACCGACCTAAGGGAGAATCGTCCCGCACCTGTAACCACA-	59
                      *   * ****  ***  * **           *   * *** *** *   *

Hu_400+        GCCCTAGGCCTCTCAGGGGTGGGTACATCAAGTGAGGAAGTCACACTGTCAGGGCAGA--	105
Mm_409+S2      ------TTCCTGATCGGTCTAGGGTCATTTCCTTCGTAGTCCTAACCACCACACCCGATT	113
                       ***    **  * **  ***    *  * *   *  **   **   * **

Hu_400+        ---------------AAGAGGGGCTGGGTGGGGCCGGACAGGCTCTGCTGGCTCAGGATC	150
Mm_409+S2      GAAGACTGGCTCCCAACCGGGAGTTGGGCATGCCACG---GCCCTCGCTGGGCCTCCCTC	170
                              *   ** * ****   * *  *   * *   *****  *    **

Hu_400+        CCGGCAGGACAGGCTGCCCAACCCAGCCCCAA-AACTCCCATCTTCTGTCCAACGCCCAA	209
Mm_409+S2      CTGTGCGCCCGGAATTTGGAGTACGCCCACGGTACCACCCGGACGCGCTAGACAGGTAAG	230
               * *   *  * *  *    *   *  ** *   * * ***     *  *  *  *   *

Hu_400+        GCCTGCTTCCTCCCTGCCCCACCTCACCTCACACTGCCCAGACTGGTGGGGCTGCATTTT	269
Mm_409+S2      GGATCCATTGAGTCCGGGCCACCGTGATGATTACTGGGCAGGAAACTCGGAGCGCAAGGG	290
               *  * * *     * *  *****         ****  ***     * **   ***

Hu_400+        T--GGGGCCGGTGCCAGGTGTGATGGAGGACAGCCCAGGTCAGGAGGTCAGGGCCCCGTT	327
Mm_409+S2      TGCGAAGCAGTTAACTGGTTTCT-----------------------GTCAGTAC--C-AT	324
               *  *  ** * *  * *** *                         *****  *  *  *

Hu_400+        TCCTCCCAAGC-CAGGATGACTCAGTGATGGACAGACTAGGCCACAAGGGGCCGAACACA	386
Mm_409+S2      TGTTGCCAAGCCAACTGTTACGCCTTGTGTGTGAGTCACCAGGGCATGTGCCAACTCAAT	384
               *  * ******  *   * ** *  **   *  ** *       ** * * *    **

Hu_400+        AAGCCAAATTAGGG-----------	400
Mm_409+S2      TAGCCGCAGTCGCGCCTCGAGCTAC	409
                ****  * * * *
```

```
Alignment #6

Hu_400+        TTCTGACACAACCTGCTACACATCTTGGATCCC--ACTTGTCAAGCCTGGCCCTAGGCCT	58
Mm_409+S3      -------TAGTCACGCGTAGCTTATTGGGTAACCTGCCTACACAGTACGAGCGTAGTAAA	53
                          *  **    * * **** *  *   * *    **   *  * ***

Hu_400+        CTCAGGGGTGGGTACATCAAGTGAGGAAGTCACACTGTCAGGGCAGAA----AGAG-GGG	113
Mm_409+S3      AATGCGGGGGGCTGA-ATACGTACCTCATTGCACCTCCGAGGCGCATGCGCGAGCGTCCC	112
                    *** ** *     * **     * *    **   ***          ** *

Hu_400+        CTGGGTGGGGCCGGACAGGCTCTGCTGGCTCAGGATCCCGGCAGGACAGGCTGCCCAACC	173
Mm_409+S3      CCGGGGGGTGCCGAACGGCATGCCATGCACCAT------TCACGTGGACACTTCCAAACC	166
               * *** ** **** ** *  *    **   **           *   *  ** ** ****

Hu_400+        CAGCCCCAAAACTCCCATCTTCTGTCCAACGCCCAAGC----CTGCTTCCTCCCTGCCCC	229
Mm_409+S3      CCG-GACTGGTATCCAATCACCGCTAGATAGCTCGGGGCGGCACGGATTCGATCTTCGCC	225
               * *   *     *** ***  *  *  *  ** *  *       *  * *   ** * **

Hu_400+        ACCTCACCTCACACTGCCCAGACTGGTGGGGCTGCATTTTTGGGGCCGGTGCCAGGTGTG	289
Mm_409+S3      GTGTCGCAAGGCCG----------AGTCGATGCTCGGGTATCACACCTAAGACTGGTGTG	275
                  ** *    *             ** *     *   * *    **   * * ******

Hu_400+        ATGGAGGACAGCCCAGGTCAG-------------GAGGTCAGGGCCCCGTTTCCTCCC-A	335
Mm_409+S3      CAGTAATACAACGGTCGTGCGCAGACCCTTTAAACGGTTCGCTGCCACGCTTCGGTTCGG	335
                 * *  *** *    **  *               * **   *** ** ***    *

Hu_400+        AGCCAGGATG----ACTCAGTGATGGACAGACTAGGCCACAAGGGGC--CGAACACAAAG	389
Mm_409+S3      GTACAGAATCGAGCTCTCTTCTCTCGCCAGCCGCAAATATCACCCGCGTCACACGTTTTA	395
                  *** **      ***     * * *** *      *  *   **  *  **

Hu_400+        CCAAATTAGGG---	400
Mm_409+S3      CCAGATTGACGCAC	409
```

*** ***	*

**Alignment #7.** Shown below is the pair-wise alignment of the core human *ppl* enhancer (Hu_400+) aligned to the 467 bp enhancer block from zebrafish (Zf_467+). The 467 bp zebrafish sequence corresponds to the enhancer fragment with the highest similarity to the core of the human enhancer. These sequences correspond to the plus strand (+) sequences located upstream of *ppl*, which is transcribed to the left in each genome.

```
Hu_400+      TTCTGACACAACCTGCTACACATCTTGGAT-------CCCACT-TGTCAAGCCTGGCCCT	52
Zf_467+      ATCTCATTCAAAGAATGTCAAGCACTGTAGTCACCGTGCCACTAAATGAACCATGGCACT	60
              *** *  ***       **     ** *         *****   * ** * **** **

Hu_400+      AG-GCCTCTCAGGGGTGGGTACATCAAGTGAGGAAGTCACACTGTCAGGGCAGAAAGAG-	110
Zf_467+      AAACCCTATGAAGTGATTGTTTTCTTTGTCAGGCAGTATCACTTACTCTAAAACCTGAGT	120
             *   *** * * * *   **       ** *** ***  ****  *     *    ***

Hu_400+      -GGGCTGGGTGGGGC------------CGGACA-GGCTCTGCTGGCTCAGGATCCCGGCA	156
Zf_467+      GAGGCACATTTGCAACATTCTTATTAAATGCCAGTCTTCTGCGACCTGTGACACACTTCA	180
               ***    * *                 * **    *****   **  *   * *  **

Hu_400+      GGACAGGCTGCCCAACCC----AGCCCCAAAACTCCCATCTTCTGTCCAACGCCCAAGCC	212
Zf_467+      ATACTGAGTGTGTAAGATGGGATGGTGTAGGCCTACTAGCTGATGTCTCAC---TAAATC	237
               ** *  **   **        *    *   ** * * **  ****  **    **  *

Hu_400+      TGCTTCCTCCCTGCCCCACCTCACCTCACACTGCCCAGA-----CTGGTGGGGCTGCATT	267
Zf_467+      TGTCAGATTAGTGCAGAACTGTTCCGTTCCATGCCTGGAAGTTTGTAGTGGGCCTGAAGA	297
             **     *   ***   **    **   *  ****  **      * ***** *** *

Hu_400+      TT--TGGGGCCGGTGCCAGGTGT----GATGGAGGACAGCCCAGGTCAGGAGG-------	314
Zf_467+      CCTTGATTTGCTTTGTCAGGTATGTTTAATTAAGGTTAGCGCTAAACTCTACATGACAAT	357
                       *  ** ***** *     **  ***  *** *    *   *

Hu_400+      ---------------TCAGGGCCCCGTTTCCTCCCAAGCCAGGATGACTCAGTGATGGAC	359
Zf_467+      GGTCCTCCAGGAACAGAAAGTTCACCTTCAATAGACAGCAATGCATGTTTACTTTTCTGC	417
                              * *  * * **   *    *** * *     * * *  *   *

Hu_400+      AGACTAGGC--------CACAAGGGGCCGAACACAAAGCCA-AATTAGGG	400
Zf_467+      AAAGTACCAAACCTCACCGCTAGGGGGAGTCGAATAGTTTTGACTGAGGC	467
             * * **           * * *****  *   *  *      * * ***
```

**Alignment #8.** Shown below is the pair-wise alignment of the core human *ppl* enhancer (Hu_400+) aligned to the reverse complement of the 467 bp block from zebrafish (Zf_409-) used in Alignment #7.

```
Hu_400+      --TTCTGAC----ACAACCTGC------TACACATCTTGGATCCCACTTGTCAAGCCTGG	48
Zf_467-      GCCTCAGTCAAAACTATTCGACTCCCCCTAGCGGTGAGGTTTGGTACTTTGCAGAAAAGT	60
                ** * *      *  *  *      **    *   *  *   ****  **     *

Hu_400+      CCCTAGGCCT----CTCA--GGGGTGGGTACATC-AAGTGAGGAAGTCACACTGTCAGGG	101
Zf_467-      AAACATGCATTGCTGTCTATTGAAGGTGAACTTTCTGTTCCTGGAGGACCATTGTCATGT	120
                 * ** *     **    *   * * ** *     *   * **   ** ***** *

Hu_400+      CAG------------------AAAGAGGGGCTGGGTGGGGCCGGACAGGCTCTGCTGGCT	143
Zf_467-      AGAGTTTAGCGCTAACCTTAATTAAACATACCTGACAAAGCAAATCAAGGTCTTCAGGCC	180
                                    * *    *  *     **    ** * *** * ***

Hu_400+      CAGGATCC-CGGCAGGACAGGCTGCCCAACCCA--GCCCCAAAACTCCCATCTTCTGTCC	200
Zf_467-      CACTACAAACTTCCAGGCATGGAACGGAACAGTTCTGCACTAATCTGACAGATTTAGTGA	240
             **  *    *  *  * ** *   *  ***       * * ** **  **  **  **

Hu_400+      AACGCCCAAGCCTGCTTCCTCCCTGCCCCACCTCACCTCACACTGCCCAGACTGGTGGGG	260
Zf_467-      GACATCAGCTAG-----TAGGCCTACACCATCCCATCTTACACACTCAGTATTGAAGTGT	295
              **  *               *** * *** * ** ** ****   *   * **  * *

Hu_400+      CTGCATTTT--------TGGGGCCGGTGCCAGGTGTGATGGAGGACAGCCCAGGTCAGGA	312
Zf_467-      GTCACAGGTCGCAGAAGACTGGCATTTAATAAGAATGTTGCAAATGTGCCTCACTCAGGT	355
              *      *           ***   *   * *  ** ** *     ***    *****

Hu_400+      GGTCAGGGCCCCGTTTCCTCCCAAGCCAGGATGACTCAGTGATGGACAGACTAGGCCACA	372
Zf_467-      TTT--------------AGAGTAAGTGATACTGCCTGACAAAGAAAACAATCACTTCATA	401
               *                   ***  *   ** ** *   *   *   *  *   ** *

Hu_400+      AGGGGCCGAACACAAAGCCAAATTAGGG--------------------------------	400
Zf_467-      GG-GTTTAGTGCCATGGTTCATTTAGTGGCACGGTGACTACAGTGCTTGACATTCTTTGA	460
              * *        **  *   * **** *

Hu_400+      -------	400
Zf_467-      ATGAGAT	467
```

**Alignment #9.** Shown below is the pair-wise alignment of the core human *ppl* enhancer (Hu_400+) aligned to the reverse sequence of the 467 bp enhancer block from zebrafish (Zf_467R). This alignment constitutes a negative control as the reverse sequence is a non-biological sequence.

```
Hu_400+      ---------TTCTGA-----------------------CACAACCTGCTACACATCT---	25
Zf_467R      CGGAGTCAGTTTTGATAAGCTGAGGGGGATCGCCACTCCAAACCATGAAACGTCTTTTCA	60
                      ** ***                       ** * * **  **   * *
                                     7-mer
Hu_400+      ------------------TGGATCCCACTTGTCAAGCCTGGCCCTAGGCCTCTCAGGGGT	67
Zf_467R      TTTGTACGTAACGACAGATAACTTCCACTTGAAAGACAAGGACCTCCTGGTAACAGTACA	120
                               *   * *******  *  *  ** ***     *  ***

Hu_400+      GGGTACATCAAGTGAGGAAGTC---------------ACACTGTCAGGGCAGAAAGAGGG	112
Zf_467R      TCTCAAATCGCGATTGGAATTAATTTGTATGGACTGTTTCGTTTAGTTCCAGAAGTCCGG	180
                 * ***  *   **** *                    * *     *****    **

Hu_400+      GCTGGGTGGGGCCGGACAGGCTCTGCTGGCTCAGGATCCCGGCAGGACAGGCTGCCCAAC	172
Zf_467R      GTGATGTTTGAAGGTCCGTACCTTGCCTTGTCAAGAC-----GTGATTAGACTGTCTAAA	235
             *    **  *   *  *   *  ***    *** **        *   ** *** * **

Hu_400+      CCAGCCCCAAAACTCCCATCTTCTGTCCAACGCCCAAGCCTGCTTCCTCCCTGCC--CCA	230
Zf_467R      TCACTCTGT----------AGTCGATCATCCGGATGTGGTAGGGTAGAATGTGTGAGTCA	285
              **  *               **  **   **     *   *  *      **     **

Hu_400+      CCTCACCTCACACTGCCCAGACTG--GTGGGGCTGCATTTTT---GGGGCCGGTGCCAGG	285
Zf_467R      TAACTTCACACAGTGTCCAGCGTCTTCTGACCGTAAATTATTCTTACAACGTTTACACGG	345
                *  * **** ** ****  *    **    *  *** **       *   * *  **

Hu_400+      TGTGATGGAGGACAGCCCAGGTCAGGAGGTCAGGGCCCCGTTTCC-TCCCAAGCCAGGAT	344
Zf_467R      AGTGAGTCCAAAA--TCTCATTCACTA--TGACGGACTGTTTCTTTTGTTAGTGAAGTAT	401
              ****      *    *    ***  *  * * ** *   **    *   *    ** **

Hu_400+      GACTCAGTGATGGACAGACTAGGCCACAAGGG---------GCCGAACACAAAGCCAAAT	395
Zf_467R      CCCAAATCACGGTACCAAGTAAATCACCGTGCCACTGATGTCACGAACTGTAAG-AAACT	460
               *  *     * **  * **   ***   *            *****   ***  ** *

Hu_400+      TAGGG--	400
Zf_467R      TACTCTA	467
             **
```

**Alignments #’s 10, 11, and 12.** Shown below are three pair-wise alignments of the core human *ppl* enhancer (Hu_400+) aligned to one of three (S1, S2, and S3) different Fisher-Yates shuffled sequences of the 467 bp plus-strand block from zebrafish (Zf_467+S1/S2/S3). These pair-wise alignments serve as negative controls.

```
Alignment #10

Hu_400+        ---------------------------------TTCTGACACAACCTGCTACACATCTTG	27
Zf_467+S1      AACCGTGTGGCATTCTTCGGTGAAACCGGTGATTTGTACCTCACTGAATCAAGCATTATA	60
                                                ** *  * **       *  ***  *

Hu_400+        GATCCCACTTGTCAAGCCTGGCC------CTAGGCCTCTCAGGGGTGGGTACAT------	75
Zf_467+S1      TAGGCAAACCCCTACGGCTAGCCCAATCAATGCGTATATCTTGAGTCTGTATTGTAATTA	120
                *  * *      * * ** ***       *  *  * **  * **  ***

Hu_400+        -CAAGTGAGGAAGTCACA--CTGTCAGGGCAGAAAGAGGGGCTGGGTGGGGCCGGACAGG	132
Zf_467+S1      ATCCTTGGGGAAGTAATCCCCGGTCAGCTCACAAAGC--TACGGGGTCTGTAATAAGTCA	178
                    ** ****** *    * *****  ** ****     * ****  *     *

Hu_400+        CTCTGCTGGCT-----------CAGGATCC----CGGCAGGACAGGCTGCCCAACCCAGC	177
Zf_467+S1      CTACGTTGCTTAGAAGCAGCTTTAGCATAGATGTAGTCTCTACAGTGTTAAAGCTCCACT	238
               **  * **  *            ** **       * *   ****  *       ***

Hu_400+        C------------CCAAAACTCCCATCTTCTGTCCAACGCCCAAGCCTGCTTCCTCCCT-	224
Zf_467+S1      CACTAGAATGATCTCAGATTATAATTCTTGTGTGTAACGTTAGCCCATCGTGTTGACCAC	298
               *             ** *       **** ***  ****      * *  *     **

Hu_400+        --GCCCCACCTCACCTCACACTGCCCAGACTGGTGGGGCTGCATTTTTGGGGCCGGTG--	280
Zf_467+S1      GCTGCACTATCCAGTCCAGACGTTTGAAATTGCTGTGCATAAGTTATATCACTCCGATAG	358
                   * *    **   ** **     * * ** ** *  *   ** *      * *

Hu_400+        CCAGGTGTGA-TGGAGGACAGCCCAGGTCAGGAGGTCAGGGCCCCGTTTCCTCCCAAGCC	339
Zf_467+S1      GGAGGTGGGTCTGTCTAACACTGAAGTTCTTGAGGTTAAA------CTTCCTG---AATC	409
                 ***** *  **    ***    ** **  ***** *         *****    *  *

Hu_400+        AGGATGACTCAGTGATGGACAGACTAGGCCACAAGGGGCCGAACACAAAGCCAAATTAGG	399
Zf_467+S1      AAATGGAATAAATACGTACACTTTTAGTACTTCGAAGTGC---TGCAACGCAGTGTTGCA	466
               *    ** * * *           ***  *      *  *     *** **    **

Hu_400+        G	400
Zf_467+S1      A	467
```

```
Alignment #11

Hu_400+        -TTCTGACACAACCT--GCTACAC---------------ATCTTGGATCCCACTTGTCAA	42
Zf_467+S2      CAAAGGAGAGAACCTTAGGAACACTTATAGCACGGGAATTTTTTGGATTCCGTTAATGGA	60
                    ** * *****  *  ****                * ****** **  *  *  *

Hu_400+        G-CCTGGCCCTAGGCCTCTCAGGGGTGGGTAC----ATCAAGT----------GAGGAAG	87
Zf_467+S2      ACTGTGCCTATACTAATATCAAACTTGATGTGATATTTAACCTTTAAAATACAGTGAAAG	120
                   ** *  **    * ***    **          * *  *          * * ***

Hu_400+        TCACACTGTCAGGGCAGAAAGAG----GGGCTGGGTGGGGCCGGACAGGCTCTGCTGGCT	143
Zf_467+S2      ATGTACTTATTATTTATATAGATTGTATGTCTCATTAACGCAAACCGTTGTCTACTTCCC	180
                   ***        * * ***      * **   *   **    *    *** **  *

Hu_400+        CAGGATCCCGGCAGGACAGGCTGCCCAACCCAGCCCCAAAACTCCCATCTTCTGTCCAAC	203
Zf_467+S2      -----TATTGGTTGACTAAGCTACACGTCTTTGCCCGT-----ACACTGTTAGGTATACA	230
                    *   **  *   * *** * *  *   ****        *  * **  **  *

Hu_400+        GCCCAAGCCTGCTTCCTCCCTGCCCCACCTCACCTCACACTGCCCAGACTGGTGGGGCTG	263
Zf_467+S2      AACCA---------------------ATTAGGTCTCCCGCTCCAGAGTAAGTGGGGGTTG	269
                 ***                     *      *** * ** *  **   *  **** **

Hu_400+        CATTTTTGGGGCCGGTGCCAGGTGTGATGGAGGACAGCCCAGGTCAGGAGGTCAGGGCCC	323
Zf_467+S2      AAATCATCGGAGCGACGCGTCATATAGTTTCTTAC---GGAGATAGGGCGCAAAGTGCAA	326
                * *  * **  **  **    * *  *     **     ** *  ** *   ** **

Hu_400+        CGTT---TCCT-----CCCAAGCCAG-------GATGACTCAGTGATGGACAGACTAGGC	368
Zf_467+S2      CTTCGATTAATATGCCGCCCATAGAACATTCAACATTACTTCGAAGCTGCCCCACCCCAC	386
               * *    *  *      ** *   *         ** ***  *     * *  **    *

Hu_400+        CACAA---------GGGGCCGA---ACACAAAGCCAAATTAGGG----------------	400
Zf_467+S2      GTGCAGGGTCTTTTGGCGGCGAGAAGGTCAACTACGTTTTAGGTAAGATTTTCTGTGACT	446
                   *         ** * ***      ***   *   *****

Hu_400+        ---------------------	400
Zf_467+S2      GTTGGTCACGCCCCGAGTCCT	467
```

```
Alignment #12

Hu_400+        ---TTCTGACACAACCTGCTACACATCTTGGATCCCACTTGT----------CAAGCCTG	47
Zf_467+S3      AGGAGTAGGCAACTAAAGATGCGTATTTTAGGTTCCTCTTCAGCCCAAACTACACACCCT	60
                      * **      * * *  ** ** * * ** ***            **  **

Hu_400+        GCCCTAGGCCTCTCAGGGGTGGGTACATCAAGTGAGGAAGTCACACTGTCAGGGCAGAAA	107
Zf_467+S3      AACGTGGGCT--TCAGGGATGTCTTTTGCATCTTACCTAATTTACACGGAGCCGCAGTAA	118
                 * * ***   ****** **  *    **  * *   * *      *     **** **

Hu_400+        GAGGGGCTGGGTGGGGCCGGACAGGCTCTGCTGGCTCAGGAT-CCCGGCAG-GACAGGCT	165
Zf_467+S3      TTAAGTATACTTTTGCAGAAGTAGTCTGCCCATAAATTGAAAGCTGGTCGCAGCTTGTTA	178
                   *  *   *  *       ** **   *       * *  *  * *   *   *
               7-mer
Hu_400+        GCCCAACCCAGCCCCAAAACTCCCATCTTCTGTCCAA--CGCCCAAGCCTGCTTCCTCCC	223
Zf_467+S3      GCCCAACTCGTATCCAGTCATTGCATCTTGTGACTTTGCAGGCTAAGCTCCCCTCATAAG	238
               ******* *    ***    *  ****** ** *      * * ****   * ** *

Hu_400+        TG----CCCCACCTCACCTCACACTGCCCAGACTGGTGGGGCTGCATTTTT-GGG-GCCG	277
Zf_467+S3      GGGTTTACTCGCTCATCCTATGCGTAATACTCATACATGGGCTTCAAGTTTAGGTAGCCC	298
                *     * * *    ***     *        *    ***** **  *** **  ***

Hu_400+        GTGCCAGGTGTG-------ATGGAGGACAGCCCAGGTCAGGAGGTCAGGGCCCCGTTTCC	330
Zf_467+S3      ATTGACGGTAGAATTACCAATTGAATATTGAGGATTAAATATGGCACGGGGACCTAGACG	358
                *    ***          ** **  *  *   *    *   **   ***  **    *

Hu_400+        TCCCAA-GCCAGGATGACTCAGTGATGGACAGACTAGGCCACAAGGGGCCGAACACAAAG	389
Zf_467+S3      TCACAAAGTCTTCGTTAATCT-AGGTGTACAATCTTTGTACTCATGTGGCGTCCGAAAGG	417
               ** *** * *    * * **   * ** ***  **  *     * * * **  *  ** *

Hu_400+        CCAAATTAGGG---------------------------------------	400
Zf_467+S3      ATAAAATGAGGAAATGTTGGTCTTCAACAAACACTTCGTTCATAAGGTTT	467
                 *** *  **
```

**Alignments # 13.** For reference, shown below is a pair-wise alignment of the core human *ppl* enhancer (Hu_400+) aligned to the full-length 812 bp enhancer from the zebrafish *ppl* locus (Zf_812+).

```
Hu_400+      ------------------------------------------------------------	0
Zf_812+      CCCACCTACATTCTGGAACGGGGAGGCTCGGAGAAAAACCAGCCAATCCTGCCTGGCCCC	60


Hu_400+      ------------TTCTGACACAACCTGCTACACATCTTGGATC----------------C	32
Zf_812+      AGACATTTATTATCATGAAAAAGAGTGTTAAACATGCTTTTAAAGAAGTCATTGTGTTGT	120
                         *  *** * *   ** ** ****  *

Hu_400+      CACTTGT--------------------------------------CAAGCCTGGCCCTAG	54
Zf_812+      GACTAATAAGATGTGTTGTTAAAAGTGTCTCTTTAGGGAATCTGCCATCAAACGCTGTAA	180
              ***  *                                      **      **  **

Hu_400+      GCCTCTCAGGGGTGGGTACATCAAGTGAGGAAGTCACACTGTCAGGGCAGAAAGAGGGGC	114
Zf_812+      TTCTCCTCGGCTTG-----CTTAAGACAGAAAGGAACATTGCCTTACAGATAAG-GAGGA	234
               ***   **  **      * ***  ** ***  *** ** *        *** * **

Hu_400+      TGGGTGGGGCCGGACAGGCTCTGCTGGCTCA----GGATCCCG-GCAGGACAGGCTGCCC	169
Zf_812+      ATGATGACAGAGCAACAGTAATGCACACTGATCTACTATACTATCTGAAACATGCAGCCA	294
               * **     * *   *   ***   ** *      ** *        *** ** ***

Hu_400+      AACCCAGCCCCA-----AAACTCCCATCTTCTGTCCAACGCCCAAGC-------------	211
Zf_812+      CACCCATTCAGTCACTCTTTCTGTCATTTTCCACCTCACACCGTGATCTCATTCAAAGAA	354
              *****  *           **  *** ***   *  ** **

Hu_400+      ---------CTGCTTCCTCCCTGCCCCACC---TCACCTCACACTGCCCAGACTGGTGGG	259
Zf_812+      TGTCAAGCACTGTAGTCACCGTGCCACTAAATGAACCATGGCACTAAACCCTATGAAGTG	414
                      ***    * ** **** *         * *  ****   *    **  * *

Hu_400+      GCTGCATTTTTGGG--GCCGGTGCCAGGTGTGATGGAGGACAGCCCAGGTCAGGAGGTCA	317
Zf_812+      ATTGTTTTCTTTGTCAGGCAGTATCACTTA-----CTCTAAAACCTGAGTGAGGCACATT	469
               **  ** ** *   * * **  **  *          * * **   ** ***

Hu_400+      GGGCCCCGTTTCCTCCCAAGCCAGGATGACTCAGTGATGGACAGACTAG-----------	366
Zf_812+      TGCAACATTCTTATTAAATGCCAGTCTTCTGCGACCTGTGACACACTTCAATACTGAGTG	529
              *   *  * *  *   * *****  *    *       **** ***

Hu_400+      -G----------------------------------------------------------	367
Zf_812+      TGTAAGATGGGATGGTGTAGGCCTACTAGCTGATGTCTCACTAAATCTGTCAGATTAGTG	589
              *

Hu_400+      ------------------------------------------------------------	367
Zf_812+      CAGAACTGTTCCGTTCCATGCCTGGAAGTTTGTAGTGGGCCTGAAGACCTTGATTTGCTT	649


Hu_400+      ----------------------------------------CCACAAGGGG----------	377
Zf_812+      TGTCAGGTATGTTTAATTAAGGTTAGCGCTAAACTCTACATGACAATGGTCCTCCAGGAA	709
                                                       **** **

Hu_400+      CCGAACACAAAGCCAAATTAGGG-------------------------------------	400
Zf_812+      CAGAAAGTTCACCTTCAATAGACAGCAATGCATGTTTACTTTTCTGCAAAGTACCAAACC	769
             * ***     * *   * ***

Hu_400+      -------------------------------------------	400
Zf_812+      TCACCGCTAGGGGGAGTCGAATAGTTTTGACTGAGGCTTGGAA	812
```

**Alignments # 14.** For reference, shown below is a pair-wise alignment of the full-length human *ppl* enhancer (Hu_489+) aligned to the full-length 812 bp enhancer from the zebrafish *ppl* locus (Zf_812+). Relative to the 400 bp core enhancer used in the majority of pair-wise alignments, the full-length enhancer sequences overlaps a non-conserved Alu element (AluJr4 sub-family) on the *ppl*-proximal side and a MIR-element on the *ppl*-distal side.

```
Hu_489+      ------------------------------------------------------------	0
Zf_812+      CCCACCTACATTCTGGAACGGGGAGGCTCGGAGAAAAACCAGCCAATCCTGCCTGGCCCC	60


Hu_489+      ------------TTCTGACACAACCTGCTACACATCTTGGATC----------------C	32
Zf_812+      AGACATTTATTATCATGAAAAAGAGTGTTAAACATGCTTTTAAAGAAGTCATTGTGTTGT	120
                         *  *** * *   ** ** ****  *

Hu_489+      CACTTGT--------------------------------------CAAGCCTGGCCCTAG	54
Zf_812+      GACTAATAAGATGTGTTGTTAAAAGTGTCTCTTTAGGGAATCTGCCATCAAACGCTGTAA	180
              ***  *                                      **      **  **

Hu_489+      GCCTCTCAGGGGTGGGTACATCAAGTGAGGAAGTCACACTGTCAGGGCAGAAAGAGGGGC	114
Zf_812+      TTCTCCTCGGCTTG-----CTTAAGACAGAAAGGAACATTGCCTTACAGATAAG-GAGGA	234
               ***   **  **      * ***  ** ***  *** ** *        *** * **

Hu_489+      TGGGTGGGGCCGGACAGGCTCTGCTGGCTCA----GGATCCCG-GCAGGACAGGCTGCCC	169
Zf_812+      ATGATGACAGAGCAACAGTAATGCACACTGATCTACTATACTATCTGAAACATGCAGCCA	294
               * **     * *   *   ***   ** *      ** *        *** ** ***

Hu_489+      AACCCAGCCCCA-----AAACTCCCATCTTCTGTCCAACGCCCAAGC-------------	211
Zf_812+      CACCCATTCAGTCACTCTTTCTGTCATTTTCCACCTCACACCGTGATCTCATTCAAAGAA	354
              *****  *           **  *** ***   *  ** **

Hu_489+      ---------CTGCTTCCTCCCTGCCCCACC---TCACCTCACACTGCCCAGACTGGTGGG	259
Zf_812+      TGTCAAGCACTGTAGTCACCGTGCCACTAAATGAACCATGGCACTAAACCCTATGAAGTG	414
                      ***    * ** **** *         * *  ****   *    **  * *

Hu_489+      GCTGCATTTTTGGG--GCCGGTGCCAGGTGTGATGGAGGACAGCCCAGGTCAGGAGGTCA	317
Zf_812+      ATTGTTTTCTTTGTCAGGCAGTATCACTTA-----CTCTAAAACCTGAGTGAGGCACATT	469
               **  ** ** *   * * **  **  *          * * **   ** ***

Hu_489+      GGGCCCCGTTTCCTCCCAAGCCAGGATGACTCAGTGATGGACAGACTAG-----------	366
Zf_812+      TGCAACATTCTTATTAAATGCCAGTCTTCTGCGACCTGTGACACACTTCAATACTGAGTG	529
              *   *  * *  *   * *****  *    *       **** ***

Hu_489+      -G----------------------------------------------------------	367
Zf_812+      TGTAAGATGGGATGGTGTAGGCCTACTAGCTGATGTCTCACTAAATCTGTCAGATTAGTG	589
              *

Hu_489+      ------------------------------------------------------------	367
Zf_812+      CAGAACTGTTCCGTTCCATGCCTGGAAGTTTGTAGTGGGCCTGAAGACCTTGATTTGCTT	649


Hu_489+      ----------------------------------------CCACAAGGGG----------	377
Zf_812+      TGTCAGGTATGTTTAATTAAGGTTAGCGCTAAACTCTACATGACAATGGTCCTCCAGGAA	709
                                                       **** **

Hu_489+      CCGAACACAAAGCCAAATTAGGG-------------------------------------	400
Zf_812+      CAGAAAGTTCACCTTCAATAGACAGCAATGCATGTTTACTTTTCTGCAAAGTACCAAACC	769
             * ***     * *   * ***

Hu_489+      -------------------------------------------	400
Zf_812+      TCACCGCTAGGGGGAGTCGAATAGTTTTGACTGAGGCTTGGAA	812
```

**Alignments # 15.** For reference, shown below is a three-way multiple sequence alignment of the core human *ppl* enhancer (Hu_400+), the homologous 409 bp block from mouse *ppl* locus (Mm_409+), and the 467 bp block from the zebrafish *ppl* locus (Zf_467+), which is located in a similar position as the mammalian enhancer. Highlighted are two of the 18 conserved mammalian blocks with some similarity to zebrafish sequences.

```
Hu_400+      --TTCTGACACAACCTGCTACACAT------CTTGGATCCCACTTGTCAAGCCTGGCCCT	52
Mm_409+      --CTATCCCACAGACTGGTGACCCT-GGAG-CTCAGACACTCCACTGGAGAGCTATCCCT	56
Zf_467+      ATCTCATTCAAAGAATGTCAAGCACTGTAGTCACCGTGCCACTAAATGAACCATGGCACT	60
                *    ** *   **     *        *   *   *        *    *  * **
                                                b.  10-mer
Hu_400+      AGGCCTCTCAGG------GGTGGGTACATCAAGTGAGGAAGTCACACTGTC----AGGGC	102
Mm_409+      GGGCCTCTCAAATTCACCATGAGAGCTTCTCTGCAAGGAAGTCACCCCATC----AGGGC	112
Zf_467+      AAACCCTATGAAG-----TGATTGTTTTCTTTGTCAGGCAGTATCACTTACTCTAAAACC	115
                **                           *  *** ***  * *   *    *   *

                                                e.  9-mer            f.>>
Hu_400+      AGAAAGAGGGGCT------GGGTG----GGGCCGGACAGGCTCTGC-TGGCTCAGGATCC	151
Mm_409+      AGA--GTGGTTCT------AGGTG----GGGCTAAACAGGCTCTACCAGGCACAGTATCC	160
Zf_467+      TGAGTGAGGCACATTTGCAACATTCTTATTAAATGCCAGTCTTCTGCGACCTGTGACACA	175
              **  * **  *          *             *** **        *   *   *
             > g.>>>     h. 7-mer
Hu_400+      CGGCAGGACAGGCTGCCCAACCCA----GCCCCAAAACTCCCATCTTCTGTCCAACGCCC	207
Mm_409+      CTGCAGGGCAGACTGCCCAGCCTC----CTGTTCACG--------------CCCATGCTT	202
Zf_467+      CTTCAATACTGAGTGTGTAAGATGGGATGGTGTAGGCCTACTAGCTGATGTCTCACTAAA	235
             *  **   * *  **   *                                *  *

Hu_400+      -AAGCCTGCTTCCTCCCTGCCCC---------------ACCTCACCTCACACTGCCCAGA	251
Mm_409+      -TTTCCTGCCTGCTGCCTGCTCCTCCCCTTCTGGGCCGTGCCAGGCTCATGCAGAGCAGA	261
Zf_467+      TCTGTCAGATTAGTGCAGAACTGTTCCGTTCCATGCCTGGAAGTTTGTAGTGGGCCTGAA	295
                  * *  *  * *                                *    *     *
                        l.>>> m.>>>
Hu_400+      CTGGTGGGGCTGCATTTTTGGGGC---------------CGGTGCCAGGTGTGATGGA--	294
Mm_409+      CCAGTACAGCCGCATTCTTGGGCAA--------------GTGTCTCTGGTGTGAATGACC	307
Zf_467+      GACCTTGATTTGCTTTGTCAGGTATGTTTAATTAAGGTTAGCGCTAAACTCTACATGACA	355
                 *      ** ** *  **                           * *    **

Hu_400+      ---GGACAGCCCAGGTCAGGAGGTCAGGGCCCCGTTTCCTCCCAAGCCAGGATGACTCAG	351
Mm_409+      CTGGGGCTGGTCAGGTCAGAAGGTCAGGGCCTTGTTCCCTCCAGTGCCCAAGTGATA---	364
Zf_467+      ATGGTCC------TCCAGGAACAGAAAGTTCACCTTCAATAGACAGCAATGCATGTTTAC	409
                *  *           * *    * *  *   **   *     **

Hu_400+      TGATGGACAGACTAGGCCA-------CAAGGGG-CCGAACACAAAGCCAAATTAGGG-	400
Mm_409+      -----GATGAGTCAGGCTA-------CAAAGGGGCAGAACAAGAAACCAAACCGAGT-	409
Zf_467+      TTTTCTGCAAAGTACCAAACCTCACCGCTAGGGGGAGTCGAATAGTTTTGACTGAGGC	467
                          *    *           ***   *   *  *      *    *

*	  96 count (3-way identical characters)
-	 158 count (null characters)
```

**Alignments # 16.** For reference, shown below is a three-way multiple sequence alignment of the core human *ppl* enhancer (Hu_400+), the homologous 409 bp block from mouse *ppl* locus (Mm_409+), and the reverse complement of the 467 bp block from the zebrafish *ppl* locus (Zf_467-). This alignment serves as a control alignment for Alignment #15, which features far fewer insertions of null characters (158 dashes versus 227 dashes). However, the number of alignment columns with 3-way identity is similar in both (96 and 97 * columns).

```
Hu_400+      ------TTCTGACACAACCTGCTACACAT----CTTGGATC--CCACTTGTCAAGCCTGG	48
Mm_409+      ------CTATCCCACAGACTGGTGACCCTGGAGCTCAGACA--CTCCACTGGAGAGCTAT	52
Zf_467-      GCCTCAGTCAAAACTATTCGACTCCCCCTAGCGGTGAGGTTTGGTACTTTGCAGAAAAGT	60
                    *       *  *   *   * *     *  *        *     *
                  a.  9-mer                                        c. 10-mer
Hu_400+      CCCTAGGCCTCTCAGG------GGTGGGTACATCAAGTG-AGGAAGTCACACTGTCAGGG	101
Mm_409+      CCCTGGGCCTCTCAAATTCACCATGAGAGCTTCTCTGCA-AGGAAGTCACCCCATCAGGG	111
Zf_467-      AAACATGCATTGCTGTCTATTGAAGGTGAACTTTCTGTTCCTGGAGGACCATTGTCAT-G	119
                   ** *  *                       *     * **   *    ***  *
                                                          e.  9-mer
Hu_400+      CAGAAAGAGGGGCT-------------------GGGTGGGGCCGGACAGGCTCTGC-TGG	141
Mm_409+      CAGA--GTGGTTCT-------------------AGGTGGGGCTAAACAGGCTCTACCAGG	150
Zf_467-      TAGAGTTTAGCGCTAACCTTAATTAAACATACCTGACAAAGCAAATCAAGGTCTTCAGGC	179
              ***     *  **                    *     **    ** * *** *  *

Hu_400+      CTCAGGATCCCGGCAGGACAGGCTGCCCAACC--CAGCCCCAAAACTCCCA--TCTTCTG	197
Mm_409+      CACAGTATCCCTGCAGGGCAGACTGCCCAGCC--TCCTGTTCACG---------------	193
Zf_467-      CCACTACAAACTTCCAGGCATGGAACGGAACAGTTCTGCACTAATCTGACAGATTTAGTG	239
             *         *  *  * **     *  * *           *

Hu_400+      TCCAACGCCCAAGCCTGCTTCCTCCCTGCCCC---------------ACCTCACCTCACA	242
Mm_409+      -CCCATGCTTTTTCCTGCCTGCTGCCTGCTCCTCCCCTTCTGGGCCGTGCCAGGCTCATG	252
Zf_467-      AGACATCAGCTAGTAGGCC-------TACACCATCCCATCTTACACACTCAGTATTGAAG	292
                 *           **        * * **                 *     * *
                                                          n. 7-mer
Hu_400+      C---TGCCCAGACTGGTGGGGCTGCATTTTTGGGGC-CGGTGCCAGGTGTGATGGA-----	294
Mm_409+      C---AGAGCAGACCAGTACAGCCGCATTCTTGGGCAAGTGTCTCTGGTGTGAATGA-----	305
Zf_467-      TGTGTCACAGGTCGCAGAAGACTGGCATTTAATAAGAATGTTGCAAATGTGCCTCACTCAG	353
                       * *        * *   * *         **  *   ****    *

                                                              p.   11-mer
Hu_400+      ---------------GGACAGCCCAGGTCAGG-----------------AGGTCAGGGCC	322
Mm_409+      ----------CCCTGGGGCTGGTCAGGTCAGA-----------------AGGTCAGGGCC	338
Zf_467-      GTTTTAGAGTAAGTGATACTGCCTGACAAAGAAAACAATCACTTCATAGGGTTTAGTGCC	413
                               * *        **                   * * ** ***

Hu_400+      CCGTTTCCTCCCAAGCCAGGATG-ACTCAGTGATGGACAGACTAGGCCACAAGGGG-CCG	379
Mm_409+      TTGTTCCCTCCAGTGCCCAAGTG-ATA--------GATGAGTCAGGCTACAAAGGGGCAG	388
Zf_467-      ATGGTTCATTTAGTGGCACGGTGACTACAGTGCTTGACATTCTTTGAATGA---------	464
               * * * *     * *    **            **        *    *

Hu_400+      AACACAAAGCCAAATTAGGG	400
Mm_409+      AACAAGAAACCAAACCGAGT	409
Zf_467-      ----------------GAT-	467


*  97 count (3-way identical characters)
- 227 count (null characters)
```

**Alignments # 17.** For reference, shown below is a three-way multiple sequence alignment of the core human *ppl* enhancer (Hu_400+), the homologous 409 bp block from mouse *ppl* locus (Mm_409+), and the (non-biological) reverse sequence of the 467 bp block from the zebrafish *ppl* locus (Zf_467R). This alignment serves as a control alignment for Alignment’s #15 and #16, which features far fewer insertions of null characters (158 dashes versus 227 dashes). While the number of alignment columns with 3-way identity is similar (99 * columns versus 96 and 97 * columns in Alignments #15 and #16, respectively), Alignment #16 features many more insertions of null characters (248 dashes inserted versus 158 and 227 dashes in Alignments #15 and #16, respectively).

```
Hu_400+      -------TTCTGACACAACCTGCTACACAT------------CTTGGATCCCACTTGTCA	41
Mm_409+      -------CTATCCCACAGACTGGTGACCCTGG--------AGCTCAGACACTCCACTGGA	45
Zf_467R      CGGAGTCAGTTTTGATAAGCTGAGGGGGATCGCCACTCCAAACCATGAAACGTCTTTTCA	60
                       *   * *  ***       *            *   **  *  *     *

Hu_400+      AGCCTGGCCCTAGGCCTCT-----CAGG------GGTGGGTACATCAAGTGAGGAAGTCA	90
Mm_409+      GAGCTATCCCTGGGCCTCT-----CAAATTCACCATGAGAGCTTCTCTGCAAGGAAGTCA	100
Zf_467R      TTTGTACGTAACGACAGATAACTTCCACTTGAAAGACAAGGACCTCCTGGTAACAGTACA	120
                 *       * *   *     *                       *  *  *   **

Hu_400+      CACTGTCAGGGCAGAAAGAGGGGCTGGGTGGGGCCGGACAGGCTCTGC-TGGCTCAGGAT	149
Mm_409+      CCCCATCAGGGCAGA--GTGGTTCTAGGTGGGGCTAAACAGGCTCTACCAGGCACAGTAT	158
Zf_467R      T-CTCAAATCGCG----ATTGGAATTAATTTGTATGGACTGTTTCGTTTAGTTCCAGAAG	175
               *    *  **        *   *   *  *     ** *  **     *   *** *

Hu_400+      CCCGGCA------GGACAGGCTGCCCAACCCAGCCCCAAAACTCCCATCTTCTGTCCAAC	203
Mm_409+      CCCTGCA------GGGCAGACTGCCCAGCCTCCTGTTCACG--------------CCCAT	198
Zf_467R      TCCGGGTGATGTTTGAAGGTCCGTACCTTGCCTTGTCAAGACGTGATTAGACTGTCTAAA	235
              ** *         *   * * *  *            *                *  *

Hu_400+      GCCCAAGCCTGCTTCCTCCCTGCCCC-----------------------------ACCTC	234
Mm_409+      GCTTTTTCCTGCCTGCTGCCTGCTCCTCCCCTTCTGGGCCG--------------TGCCA	244
Zf_467R      TCACT-----CTGTAGT---CGATCATCCGGATGTGGTAGGGTAGAATGTGTGAGTCATA	287
              *           *  *    *  *

Hu_400+      ACCTCACA--CTGCCCAGACTG--GTGGGGCTGCATTTTTGGGGC-CGGTGCCAGGTGTG	289
Mm_409+      GGCTCATG--CAGAGCAGACCA--GTACAGCCGCATTCTTGGGCAAGTGTCTCTGGTGTG	300
Zf_467R      ACTTCACACAGTGTCCAGCGTCTTCTGACCGTAAATTATTCTTACAACGTTTA-------	340
                ***      *  ***       *        *** **        **

Hu_400+      ATGGA-----GGACAGCCCAGGTCAGGAGGTCAGGGCCCCGTTTCCTCCCAAGCCAGGAT	344
Mm_409+      AATGACCCTGGGGCTGGTCAGGTCAGAAGGTCAGGGCCTTGTTCCCTCCAGTGCCCAAGT	360
Zf_467R      ------------------CACGGAGTGAGTCCAAAATCTCATTCACTATGACGGA---CT	379
                               ** *     **  **    *   **  **     *      *

Hu_400+      GACTCAGTGATGGACAGACTAGGCCACAAGG-----------------------------	375
Mm_409+      GATA--------GATGAGTCAGGCTACAAAG-----------------------------	383
Zf_467R      GTTTCTTTTGTTAGTGAAGTATCCCAAATCACGGTACCAAGTAAATCACCGTGCCACTGA	439
             *                   *  * * *

Hu_400+      -GG-CCGAACACAAAGCCAAATTAGGG-	400
Mm_409+      -GGGCAGAACAAGAAACCAAACCGAGT-	409
Zf_467R      TGTCACGAACTGTAAGAAACTTACTCTA	467
              *    ****   **   *

*  99 count (3-way identical characters)
- 248 count (null characters)
```

**Alignments #’s 18, 19, and 20.** Shown below are three 3-way alignments of the core mammalian *ppl* enhancers (Hu_400+ and Mm_409+) aligned to one of three (S1, S2, and S3) different Fisher-Yates shuffled sequences of the 467 bp plus-strand block from zebrafish (Zf_467+S1/S2/S3). These multiple-sequence alignments serve as negative controls.

```
Alignment #18

Hu_400+        ----TTCTGACACAACCTGCTACACAT----CTTGGATCCC-------------ACTTGT	39
Mm_409+        ----CTATCCCACAGACTGGTGACCCTGGAGCTCAGACACT-------------CCACTG	43
Zf_467+S1      AACCGTGTGGCATTCTTCGGTGAAACCGGTGATTTGTACCTCACTGAATCAAGCATTATA	60
                    * *  **      * *           *  *   *

Hu_400+        CAAGCCTGGCCCTAGGCCTCTCAGG-------------------------GGTGGGTACA	74
Mm_409+        GAGAGCTATCCCTGGGCCTCTCAAATTCAC-------------------CATGAGAGCTT	84
Zf_467+S1      TAGGCAAACCCCTACGGCTAGCCCAATCAATGCGTATATCTTGAGTCTGTATTGTAATTA	120
                *       ****  * **  *

Hu_400+        TCAAGTGAGGAAGTCACACTGTCAGGGCAGAAAGAGGGGCTGGGTGGGGCCGGACAGGCT	134
Mm_409+        CTCTGCAAGGAAGTCACCCCATCAGGGCAGA--GTGGTTCTAGGTGGGGCTAAACAGGCT	142
Zf_467+S1      ATCCTTGGGGAAGTAATCCCCGGTCAGCTCACAAAGCTACGGGGTCTGT--AATAAGTCA	178
                       ****** *  *       **  *    *   *  ***  *       ** *

Hu_400+        CTGC-TGGCTCAGGATCCCGGCAGGACAGGCTG--CCCAACCCAGCCCCAAAACTCCCAT	191
Mm_409+        CTACCAGGCACAGTATCCCTGCAGGGCAGACTG--CCCAGCCTCCTGTTCACG-------	193
Zf_467+S1      CTACGTTGCTTAGAAGCAGCTTTAGCATAGATGTAGTCTCTACAGTGTTAAAGCTCCACT	238
               ** *   **  ** * *       *      **    *            *

Hu_400+        C-------TTCTGTCCAACGCCCAAGCCTGCTTCCTCCCTGCCCC---------------	229
Mm_409+        --------------CCCATGCTTTTTCCTGCCTGCTGCCTGCTCCTCCCCTTCTGGGCCG	239
Zf_467+S1      CACTAGAATGATCTCAGATTATAATTCTTGTGTGTAACGTTAGCCCATCGTG-TTGACCA	297
                             *  *        * **  *    * *   **

Hu_400+        ACCTCACCTCACACTGCCCAGACTGGTGGGGCTGCATTTTTGGGGC-CGGT---GCCAGG	285
Mm_409+        TGCCAGGCTCATGCAGAGCAGACCAGTACAGCCGCATTCTTGGGCAAGTGT---CTCTGG	296
Zf_467+S1      CG-CTGCACTATCCAGTCCAGACGTTTGAA-----ATTGCTGTGCATAAGTTATATCACT	351
                         *  * *  *****   *        ***  ** *     **     *

Hu_400+        TGTGATGGA-----GGACAGCCCAGGTCAGGAGGTCAGGGCCCCGTTTCCTCCCAAGCCA	340
Mm_409+        TGTGAATGACCCTGGGGCTGGTCAGGTCAGAAGGTCAGGGCCTTGTTCCCTCCAGTGCCC	356
Zf_467+S1      CCGATAGGGAGGTGGGTCTGTCTAACACTGAAGTTCTTGAGGTTAAA-CTTCCTGAATCA	410
                      *      ** * *   *   * * ** **  *         * ***     *

Hu_400+        GGATGACTCA-GTGATGGACAGACTAGGCCACAAGGGG-CCGAACACAAAGCCAAATTAG	398
Mm_409+        AAGTGATA---------GATGAGTCAGGCTACAAAGGGGCAGAACAAGAAACCAAACCGA	407
Zf_467+S1      AATGGAATAAATACGTACACTTTTAGTACTTCGAAGTGCTGCAACGCAGTGTTGCAA---	467
                   **            *         *  * * * *    ***          *

Hu_400+        GG	400
Mm_409+        GT	409
Zf_467+S1      --	467

* 110 count (3-way identical characters)
- 170 count (null characters)
```

```
Alignment #19

Hu_400+        ----------TTCTGACACAACCTGCTACACAT----CTT---GGAT-CCCACTTGTCAA	42
Mm_409+        ----------CTATCCCACAGACTGGTGACCCTGGAGCTC---AGAC-ACTCCACTGGAG	46
Zf_467+S2      AAAGGAGAGAACCTTAGGAACACTTATAGCACGGGAATTTTTTGGATTCCGTTAATGGAA	60
                            *     *  **  *           *     **   *        *

Hu_400+        GCCTGGCCCTAGGCCTCTCAGG------------------GGTGGGTACATCAAGTGAGG	84
Mm_409+        AGCTATCCCTGGGCCTCTCAAATT------------CACCATGAGAGCTTCTCTGCAAGG	94
Zf_467+S2      CTGTGCCTATACTAATATCAAACTTGATGTGATATTTAACCTTTAAAATACAGTGAAAGA	120
                  *  *  *     * ***                                  *  **

Hu_400+        AAGTCACACTGTCAGGGCAGAAAGAGGGGCTGGGTGGGGCCGGACAGGCTCTGC-TGGCT	143
Mm_409+        AAGTCACCCCATCAGGGCAGA--GTGGTTCTAGGTGGGGCTAAACAGGCTCTACCAGGCA	152
Zf_467+S2      TGTACTTATTATTTATATAGATTGTATGTCTCATTAACGCAAACCGTTGTCTACTTCCCT	180
                   *      *      ***  *     **   *   **    *    *** *    *

Hu_400+        CAGGATCCCGGCAGGACAGGCTGCCCAACCCAGCCCCAAAACTCCCATCTTCTGTCCAAC	203
Mm_409+        CAGTATCCCTGCAGGGCAGACTGCCCAGCCTCCTGTTCACG--------------CCCAT	198
Zf_467+S2      A-----------TTGGTTGACTAAGCTACACGTCTTTGCCCGTACACTGTTAGGTATACA	229
                             *   * **   *  *

Hu_400+        GCCCAAGCCTGCTTCCTCCCTGCCCC---------------ACCTCACCTCACACTGCCC	248
Mm_409+        GCTTTTTCCTGCCTGCTGCCTGCTCCTCCCCTTCTGGGCCGTGCCAGGCTCATGCAGAGC	258
Zf_467+S2      A-------------ACCAATTAGGTCTCCCGCTCCAGAGTAAGTGGGGGTTGAAATCATC	276
                              *    *    *                       *         *

Hu_400+        AGACTG----GTGGGGCTGCATTTTTGG--GGC-CGGTGCCAGGTGTGATGGA-----GG	296
Mm_409+        AGACCA----GTACAGCCGCATTCTTGG--GCAAGTGTCTCTGGTGTGAATGACCCTGGG	312
Zf_467+S2      GGAGCGACGCGTCATATAGTTTCTTACGGAGATAGGGCGCAAAGTGCAACTTCGATT-AA	335
                **       **      *  *  *  *  *     *      ***  *

Hu_400+        ACAGCCCAGGTCAGGAGGTCAGGGCCCCGTTTCCTCCCAAGCCAGGATG-ACTCAGTGAT	355
Mm_409+        GCTGGTCAGGTCAGAAGGTCAGGGCCTTGTTCCCTCCAGTGCCCAAGTG-ATA-------	364
Zf_467+S2      TATGCCGCCCATAGAACATTCA----ACATT-ACTTCGAAGCTGCCCCACCCCACGTGCA	390
                  *        ** *  *          **  ** *   **

Hu_400+        GGACAGACTAGGCCACAAGGGG-CCGAACACAAA---GCCA-AAT--TAGGG--------	400
Mm_409+        -GATGAGTCAGGCTACAAAGGGGCAGAACAAGAA---ACCA-AAC--CGAGT--------	409
Zf_467+S2      GGGTC-TTTTGGCGGCGAGAAGGTCAACTACGTTTTAGGTAAGATTTTCTGTGACTGTTG	449
                *        ***  * *   *    *  *          *  *      *

Hu_400+        -----------------	400
Mm_409+        -----------------	409
Zf_467+S2      GTCACGCCCCGAGTCCT	466

*  86 count (3-way identical characters)
- 216 count (null characters)
```

```
Alignment #20

Hu_400+        TTCTGACACAACCTGCTACACAT----CTTGGATCCCACTTGTCAAGCCTGGCCCTAG--	54
Mm_409+        CTATCCCACAGACTGGTGACCCTGGAGCTCAGACACTCCACTGGAGAGCTATCCCTGG--	58
Zf_467+S3      ---AGGAGTAGGCAACTAAAGATGCGTATTTTAGGTTCCTCTTCAGCCCAAACTACACAC	57
                        *  *   *     *     *   *     *     *   *   *

Hu_400+        --GCCTCTCAGG------GGTGGGTACATCAAGTGAGGAAGTCACACTGTCAGGGCAGAA	106
Mm_409+        --GCCTCTCAAATTCACCATGAGAGCTTCTCTGCAAGGAAGTCACCCCATCAGGGCAGA-	115
Zf_467+S3      CCTAACGTGGGCTTCAGGGATGTCTTTTGCATCTTACCTAATTTACACGGAGCCGCAGTA	117
                      *                           *   * *            ****

Hu_400+        AGAGGGGCTGGGTGGGGCCGGACAGGCTCTGC-TGGCTCAGGATCCCGGCAGGACAGGCT	165
Mm_409+        -GTGGTTCTAGGTGGGGCTAAACAGGCTCTACCAGGCACAGTATCCCTGCAGGGCAGACT	174
Zf_467+S3      ATTAAGTATACTTTTGCAGAAGTAGTCTGCCCATAAATTGAAAGCTGGTCGCAGCTTGTT	177
                       *   *  *       ** **   *          * *    *    *    *

Hu_400+        GCCCAACCCAGC-CCCAAAACTCCCATCTTCTGTCCAA--CGCCCAAGCCTGCTTCCTCC	222
Mm_409+        GCCCAGCCTCCT-GTTCACG--------------CCCA--TGCTTTTTCCTGCCTGCTGC	217
Zf_467+S3      AGCCCAACTCGTATCCAGTCATTGCATCTTGTGACTTTGCAGGCTAAGCTCCCCTCATAA	237
                 **   *                          *      *      *   * *  *

Hu_400+        CTGCCC-C---------------ACCTCACCTCACACTGCCCAGACTGGTGGGGCTG---	263
Mm_409+        CTGCTC-CTCCCCTTCTGGGCCGTGCCAGGCTCATGCAGAGCAGACCAGTACAGCCG---	273
Zf_467+S3      GGGGTTTACTCGCTCATCCTATGCGTAATACTCATACATGGGCTTCAAGTTTAGGTAGCC	297
                 *                           ****  *        *  **   *

Hu_400+        CATTTTTGGGGC-------------CGGTGCCAGGTGTGATGGA-----GGACAGCCCAG	305
Mm_409+        CATTCTTGGGCA------------AGTGTCTCTGGTGTGAATGACCCTGGGGCTGGTCAG	321
Zf_467+S3      CATTGACGGTAGAATTACCAATTGAATATTGAGGATTAAATATGGCACGGGGACCTAGAC	357
               ****   **                   *    * *   *         **       *

Hu_400+        GTCAGGAGGTCAGGGCCCCGTTTCCTCCCAAGCCAGGATGACTCAGTGATGGACAGACTA	365
Mm_409+        GTCAGAAGGTCAGGGCCTTGTTCCCTCCAGTGCCCAAGTGATA--------GATGAGTCA	373
Zf_467+S3      GTCACAAAGTCTTCGTTAATCTAGGTGTACAATCTTTGTACTCATGTG--GCGTCCGAAA	415
               ****  * ***   *      *   *       *    *                    *

Hu_400+        GGCCACAAGGGG-CCG-----AACACAAAGCCAAATTAGGG-----------	400
Mm_409+        GGCTACAAAGGGGCAG-----AACAAGAAACCAAACCGAGT-----------	409
Zf_467+S3      GGATAAAATGAGGAAATGTTGGTCTTCAACAAACACTTCGTTCATAAGGTTT	467
               **  * ** * *           *   **   * *    *

*  90 count (3-way identical characters)
- 140 count (null characters)
```
